# MetaNetMap: automatic mapping of metabolomic data onto metabolic networks

**DOI:** 10.64898/2026.01.05.697697

**Authors:** Coralie Muller, Juliette Audemard, Sylvain Prigent, Clémence Frioux

## Abstract

**Motivation:** Metabolic networks represent genome-derived information about the biochemical reactions that cells are capable of performing. Mapping omic data onto these networks is important to refine model simulations. However, metabolomic data mapping remains very challenging due to difficulties in identifier reconciliation between annotation profiles and metabolic networks.

**Results:** MetaNetMap is a Python package designed to automatise the process of mapping metabolomic data onto metabolic networks. It includes several layers of identifier matching, the use of customisable databases, and molecular ontology integration to suggest the most matches between experimentally-identified metabolites and molecules defined in the network. We demonstrate its usability and the quality of automated mapping using two datasets.

**Availability and Implementation:** MetaNetMap is an open source python package and publicly available under the GPLv3 licence. Source code is freely available on GitHub: https://github.com/coraliemuller/metanetmap. Data and code used in the application cases of this paper can be found at: https://doi.org/10.57745/ESFLR8.

## 1 Introduction

Integration of multi-omic data is crucial for generating high-quality computational models in systems biology. Genomics offers insights into the potential metabolic activities of an organism. Genome-scale metabolic networks (GSMNs) gather such information as a collection of metabolic reactions supported by gene-protein-reaction relationships inferred from the genome [1]. In contrast, metabolomics provides an overview of the molecules produced by a biological system under defined environmental conditions. Annotation profiles derived from metabolomic data summarise the outcome of metabolite identification obtained by matching raw signals to reference databases.

Metabolic models, which rely on GSMN structure to predict cellular behaviour in response to environmental conditions, benefit from the integration of multi-omic data, such as transcriptomics for expressed genes. Including metabolomics in such models has multiple interests: i) identifying missing components in GSMNs [2], ii) evaluating the quality of the reconstruction, iii) validating and/or constraining model predictions with observations. Mapping identified molecules from metabolomic experiments onto the structure of the metabolic network is attractive, both for the study of individual populations or communities of microorganisms, and for the characterisation of both primary and specialised metabolism.

Challenges associated with such mapping hinder the systematic integration of the data, often necessitating manual or semi-manual mapping. This process can be time-consuming, burdensome and hardly reproducible for large datasets. Another source of complexity is the presence of inconsistencies in the source data, for instance through non-standard structures such as for SMILES or InChI. Additionally, metabolite annotations from metabolomic profiles are often partial, incomplete, or inconsistently represented. The absence of standardised identification across studies leads to tailored, study-specific mapping efforts and highlights the need for a more streamlined and automated process. Existing efforts in database-to-database mapping, such as MetaNetX [3], provide an important foundation for mapping. However, the dispersion of information across various knowledge bases and input files necessitates additional development efforts to ensure that the mapping process fully leverages both the available resources and the metadata associated with input metabolomics and GSMNs.

In this paper, we present MetaNetMap, a Python package designed to overcome these difficulties.

MetaNetMap automatically matches metabolite information between metabolomic data and GSMNs. It leverages direct mapping taking into account metadata of input files, and indirect mapping by exploiting conversion data tables built from third-party knowledge bases. It additionally considers partial matching techniques to improve mapping rates.

## 2 Overview of the method

### General description

MetaNetMap aims at automatically mapping metabolites identified from metabolomic data processing to those described in a single GSMN or a collection of them (*community* mode). The general process is described in Figure 1. MetNetMap performs two main steps: i) direct matching using inputs’ metadata, and ii) indirect matching with an external resource to bridge identifiers. An optional step relies on partial matching to increase the number of mapped metabolites. The tool uses metadata extracted from the GSMN and, when available, from annotation profiles. It can also incorporate a user-provided database to match metabolite identifiers across multiple knowledge sources.

**Figure 1.**
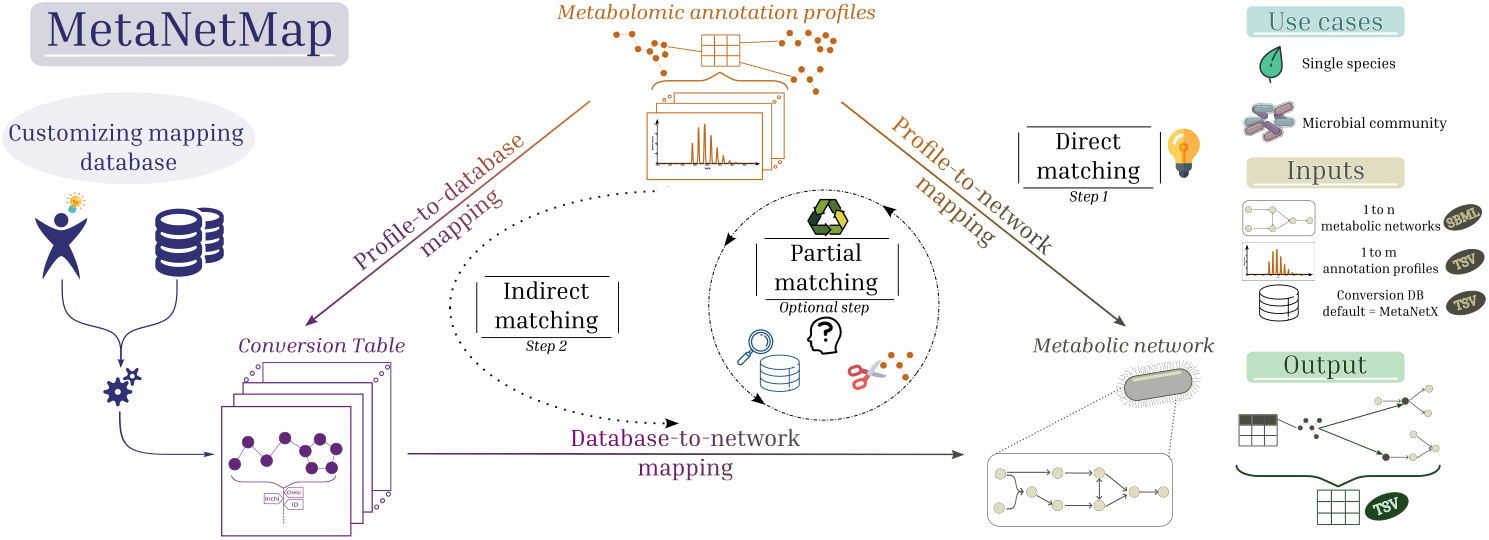
Overview of MetaNetMap process. MetaNetMap provides both direct and indirect mapping between metabolomic annotation profiles and GSMNs. Indirect mapping relies on a conversion table that can be created by the tool using MetaCyc or MetaNetX third-party knowledge bases. Each mapping procedure includes first direct, then indirect mapping. Optionally, partial matching can be performed to rescue additional mapping by relaxing matching constraints.

MetaNetMap additionally constructs the conversion table used for indirect mapping using third-party databases (MetaNetX, MetaCyc or custom ones).

### Mapping database construction

Indirect mapping necessitates a conversion table composed of metabolite metadata that can bridge identifiers from metabolomic inputs to GSMNs. MetaNetMap can construct this input. Current implementation supports the use of MetaCyc [4] and MetaNetX [3] knowledge bases. In this mode, MetaNetMap combines the structured information extracted from Meta-Cyc’s ‘compounds.dat’ or from MetaNetX’s ‘chem xref.tsv’ and ‘chem prop.tsv’ files. Those files contain metabolite information and cross references in the corresponding database (identifiers, chemical structures, molecular formulas, etc.). During this step, it is also possible to incorporate custom identifiers or additional information, for instance resulting from prior manual mapping efforts. The resulting unified table acts as a comprehensive knowledge base, enabling the tool to search among all known identifiers for a given metabolite and match them between metabolomic data and GSMNs. More details about this step are provided in Supplementary.

### Mapping modes

Mapping is the core of MetaNetMap. Two modes are available:

- The *Classic mode* maps one or several annotation profiles onto a single GSMN.
- The *Community mode* maps one or several annotation profiles onto multiple GSMNs. This mode is useful when processing metabolomic data obtained from a microbial community where each population is associated to one GSMN.

For both modes, data is pre-processed and the following steps are performed sequentially during the mapping procedure: i) matching GSMN metadata with annotation profiles (direct mapping), ii) matching metabolomic data with the conversion table, and iii) matching conversion table’s metabolites with GSMN’s metabolite metadata (Fig. 1).

### Partial matches

It is possible that neither direct mapping (via inputs’ metadata) nor indirect one (through the conversion table) succeed at matching metabolites between annotation profiles and GSMNs. An option was developed to retrieve additional mapping cases by considering partial match of certain identifiers. It is a post-processing step applied to identifiers that were not successfully mapped during the initial mapping procedure. These unmatched entries are re-evaluated using specific strategies, which increase the chances of finding a match (e.g., via ChEBI, InChIKey, or enantiomer simplification). More details are available in Supplementary.

### Handling Ambiguities

Using a large amount of data increases the number of matches for certain entries and, consequently, the risk of ambiguity where a metabolite matches to several entries of the conversion datatable, metabolomic data, or GSMN. Those multiple matches are captured by MetaNetMap and their resolution is left to the user. Three main types of ambiguities can be mentioned:

- One metabolite of the metabolomic annotation profile maps onto two or more distinct metabolites of the GSMN.
- One metabolite of a metabolomic profile maps onto two or more metabolites in the conversion table.
- Two distinct metabolites of the metabolomic profiles map onto the same metabolite of the GSMN.

These cases are illustrated in greater detail in the Supplementary Materials, with examples provided from the application datasets.

### Outputs

Resulting mapping information after MetaNetMap processing is provided in tabular format, where each line describes a match between the metabolomic data and the conversion table and/or the GSMN(s), or an absence of match. For each match, mapped identifiers and metadata are provided, and the GMSN(s) in which the metabolite was found for community mode is (are) listed. Matching steps and their outcomes are further described in a log file dedicated to tracing the mapping procedure, ensuring reproducibility of results.

## 3 Mapping procedure

The mapping procedure consists of three steps connecting respectively i) metabolomic profiles to GSMNs, metabolomic profiles to the reference database, and iii) reference database to GSMNs (Fig. 1). Preprocessing is described in Supplementary.

- *Step 1 - Direct mapping from annotation profiles to GSMNs*. This step tests matching metabolite identifiers, and associated metadata, of annotation profiles to metabolites of GSMNs and their own metadata described in the SBML file. The resulting matches are nonetheless searched in the reference database to provide more comprehensive metadata for those metabolites and to identify possible ambiguous matches when the mapped identifier is not unique to a metabolite.
- *Step 2 - Matching annotation profiles and the reference database*. In this first stage of the indirect mapping, metabolite identifiers and their metadata, that were not matched in Step 1, are compared to those of the conversion table. Duplicate checks are performed because several identifiers are tested for a metabolite and those could be associated to distinct entries of the reference database (See Supplementary).
- *Step 3 - Matching metabolites of the reference database and GSMN compound metadata*. This is the second stage of indirect mapping, for metabolomic entries that were successfully associated to the reference database at Step 2. All identifiers of these metabolites are tested against GSMN information. Unsuccessful mapping for those metabolites will still be notified to users and be part of MetaNetMap’s output.

## 4 Application

We demonstrate MetaNetMap’s relevance using two previously published case studies that involved mapping metabolomic data onto GSMNs. The first one is focused on the identification of metabolites associated with Parkinson’s Disease (PD)[5]. From the analysis of 87 human metabolomic studies, the authors compiled a list of 928 metabolites that were mapped onto a version of the human metabolic model Recon3 3D [6] called Recon3 3. The second study used integrated metabolomics and network pharmacology to investigate the cardioprotective effects of myricetin after a week of high-intensity exercise in mice [7].

In both studies, no indication of the mapping procedure was available, suggesting that in-house automatic and/or manual tailored mapping was performed. We used MetaNetMap (v1.1.0) to automatise the mapping and checked the resulting metabolite matches with the corresponding studies’ results. We also assessed the impact of the third-party reference knowledge base (MetaCyc or MetaNetX) used for reference database construction on the mapping rates. Results are described in Table 1 and details of methods, as well as additional results, are provided in Supplementary.

**Table 1:**
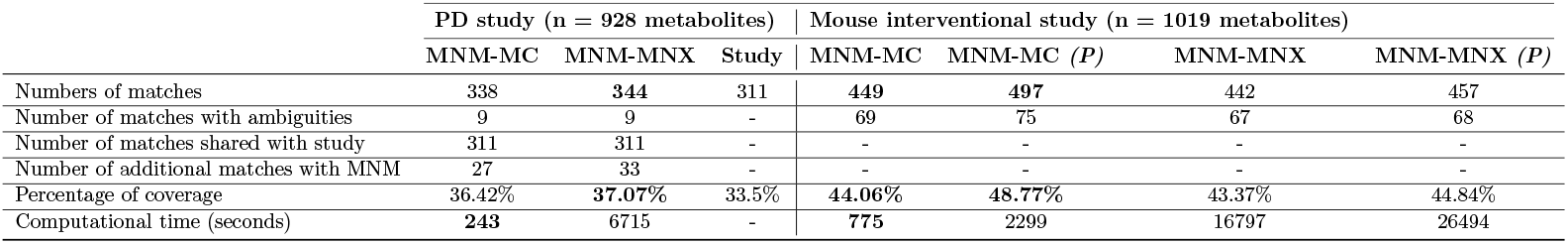
Comparison of MetaNetMap (MNM) mapping results in the Parkinson’s disease study (“PD study”) and the mouse interventional study using Metacyc (MNM-MC) or MetaNetX (MNM-MNX) as reference database for indirect mapping. The number of matches as described in the table refers to the complete number of metabolites in the GSMN which matched with or without ambiguities. The number of matches with ambiguities refers to the number of metabolites in the GSMN with more than one match in the original metabolomic profiles. “Study” refers to the number of mapped metabolites in the original PD study. Partial matching (“*(P)*”) was applied to the Mouse study. Percentage of coverage indicates the proportion of metabolites from metabolomics that could be mapped onto the GSMN. Computational time is assessed on a personal laptop with 64Gb RAM.

## Metabolomic mapping onto the human GSMN in the context of PD

In total, 344 and 338 metabolomic annotations were successfully mapped onto the human GSMN by MetaNetMap, using MetaNetX and MetaCyc as reference databases, respectively. Of those, 311 were also identified in the corresponding study, with identical mapping outcomes. The 27 new matches that MetaCyc’s reference database identified, were also identified using MetaNetX. Those matches occurred mainly through the use of InChI and PubChem identifiers.

The additional six metabolites identified with MetaNetX relied on InChI information. These were identified with indirect mapping only because InChI identifiers from the metabolomic data did not directly correspond to those of the GSMN. The correspondence was therefore established indirectly using the reference database, by linking the metabolomic InChI identifiers to MetaNetX conversion table, and then associating the VMH identifier to the GSMN. This is a good example of the usefulness of the conversion table acting as a bridge between metabolomic data and GSMNs.

Automatic mapping yielded 9 metabolites as ambiguous among the 311 shared with the PD study. For some InChI identifiers of the metabolomic data, two metabolites were identified in the GSMN. Theoretically, this should not happen since an InChI uniquely describes a molecule, but in Recon3 3 metadata, some InChI referred to multiple metabolites which explains observed ambiguities. This study could not use partial matching, as the metabolomic data did not contain any additional metadata that could be considered.

## Metabolomic mapping onto human GSMN in the context of an interventional study (Mouse)

The second study did not permit a direct comparison of mapping results as such mapping was not performed in the original study. It was however a relevant case-study notably because annotation profiles (n=1019 metabolites) were associated to metadata that could be used for partial mapping.

MetaNetMap identified 449 metabolite matches on Recon3 3 when using MetaCyc, and 442 metabolites with MetaNetX, representing identification rates of 44.06% and 43.37%, respectively. These higher rates, compared to the previous study, associate to a number of ambiguous matches that remain and require manual verification. These ambiguities can be attributed to the use of molecular formulas during the mapping process. While formulas are advantageous when no other matches are available, their use is otherwise not recommended, as distinct metabolites can share the same formula, leading to ambiguity. The mapping rates obtained are within the expected range for this type of analysis, as the average rate of mapping between GSMNs and mass spectral information from standards is only about 40%. [2]. Obtaining such ratios with a fully automated and traceable procedure is therefore a positive outcome. Partial matching was applied on the data from this study using ChEBI identifiers were provided in annotation profiles. This resulted in an increase of matches for both Metacyc and MetaNetX, up to mapping ratios of 48.77% and 44.84%, respectively. Those partial matches are curation opportunities for the corresponding metabolites.

Overall, these two applications illustrate the relevance of MetaNetMap’s strategy for mapping metabolomic data onto GSMNs, and highlight efforts to improve mapping explainability in order to understand the origin of successful matches. Computational time of mapping is relatively low when no partial matching is performed (Table 1). The smaller size of MetaCyc database compared to MetaNetX strongly impacts the performances.

## 5 Discussion

In summary, MetaNetMap streamlines and automates the mapping of metabolites identified in metabolomic data to those described in GSMNs. This is made possible by the construction of an easy-to-use, easy-to-read intermediary conversion table, which can be quickly rebuilt using new versions of MetaNetX or MetaCyc, with the option to manually add user-defined mappings if needed. MetaNetMap covers a broader range of identifiers for the same metabolite than manual or traditional mapping approaches and offers automatic solutions when mappings are partial, providing relevant information to the user, who can then manually correct results if necessary. All resultant information is consolidated into a single, user-friendly file, with an additional log file provided for further details and explainability purposes.

## 6 Acknowledgements

We acknowledge the French National Research Agency (ANR) France 2030 PEPR Agroécologie et Numérique MISTIC ANR-22-PEAE-0011, Conseil Régional d’Aquitaine and the MetaboHUB infrastructure (MetaboHUB ANR-11-INBS-0010) for their support in this work. We acknowledge the Plafrim (see https://www.plafrim.fr) for providing the computing infrastructure. We also acknowledge Ludovic Cottret for the valuable ideas and constructive discussions that contributed to this work.

## 7 Author contributions

Conceptualisation: CM, SP, CF - Investigation: all - Software: CM - Methodology: CM, SP, CF - Validation: CM, SP, CF - Writing - Original Draft Preparation: CM, SP, CF - Writing - Review & Editing: all - Visualisation: CM, SP, CF

