## Supplementary Material for "MetaNetMap: automatic mapping of metabolomic data onto metabolic networks"

### 1 Additional information on MetaNetMap

#### 1.1 Reference database (conversion table) restrictions

The MetaCyc database file and metabolite ontology information are not included in MetaNetMap and must be provided by the user because their use requires a license. MetaNetX being freely available, it can be used by default to perform mapping. Users can supply the necessary files, or MetaNetMap can download them prior to creating the reference database. Alternatively, a custom conversion table may be provided, but it must comply with specific rules regarding column names and cannot be generated by the tool. We refer to the online documentation [1] for further details.

#### 1.2 Preprocessing of data prior mapping

##### 1.2.1 Generation of MNM IDs

An MNM_ID is assigned as a unique identifier to each row corresponding to a metabolite in the metabolomic annotation profile files provided by the user. It is generated at the beginning of the MetaNetMap run, and the profiles with the additional identifiers are written to the output directory for traceability. This enables fast and easy tracking of all IDs associated with a given metabolomic annotation and helps prevent ambiguities. For example, if two distinct metabolites from the annotation profile map to the same metabolite in the GSMN, their MNM_IDs will be different; such ambiguities are then reported in the *Partial match* column of the output, allowing users to address them more easily.

##### 1.2.2 Merging and harmonisation of data

Whether users provide single or multiple metabolomic annotation profile files, the data will be grouped according to identifier column names, such as 'UNIQUE-ID', 'COMMON-NAME', and so on. This approach allows for verification in the output file of which column each metabolite matches, and maintains a clear log file.

For metabolic networks, MetaNetMap typically extracts both the ID and the name of every metabolite. If additional information is available in the GSMNs, such as RDF metadata, the tool considers all such information for each metabolite (e.g., ChEBI, InChIKey).

During preprocessing, MetaNetMap attempts to merge metabolites of the GSMN based on their names. This is usually the most suitable criterion since names are present in all metabolic networks and are more likely to be shared directly. In contrast, the specific identifier for each metabolite, if not generated by the same automatic reconstruction tool, will differ, hindering the merging process. Additionally, metadata for each metabolic network is reviewed, attempting to match and merge entries when possible.

[1]https://MetaNetMap.readthedocs.io

During preprocessing, MetaNetMap ensures that specific columns contain the correct prefixes (e.g., "CHEBI:", "PUBCHEM:", or "INCHIKEY="). This helps prevent conflicts between numerical values in different columns and reduces the risk of mismatches by harmonising identifiers as much as possible across metabolomic data, metabolic networks, and the conversion table.

#### 1.3 Mapping details

During step 2 of the mapping procedure, it is possible that several identifiers of annotation profiles for the same metabolite may match in the conversion table. In such instances, the matches will be consolidated in the output, separated by a specific string '_AND_' in order to facilitate further parsing.

#### 1.4 Partial matching

Several strategies of partial matching can be applied depending on the identifiers that annotation profiles provide as metadata.

- ***ChEBI*** identifiers. For each unmapped metabolite associated with a ChEBI identifier, the API from EBI is used to retrieve the full ChEBI ontology [1] of the metabolite. These related terms are then remapped following the classical procedure.
- ***InChIKey***. An InChIKey is structured in multiple blocks (`XXXXXXXXXXXXXX-YYYYYYYYAB-Z`), with the first block representing the core molecular structure. To improve the likelihood of a match during Step 2 of the mapping, we extract only this primary structure. However, this approach may increase the number of matches, as multiple metabolites can share the same core structure, thus necessitating manual curation.
- ***Enantiomers***. Stereochemistry indicators (L, D, R, S) are removed. This improves matching rates, since stereochemical information is rarely specified in annotation profiles because enantiomers are generally indistinguishable by standard LC–MS methods.

After this processing step is applied to annotation profiles and metabolic network identifiers, the entire mapping pipeline is re-executed to account for the modifications. Note that, in all cases, matches based on formula are automatically designated as 'partial matches,' since the formula may be shared by several distinct metabolites.

#### 1.5 Ambiguity handling

MetaNetMap does not attempt to resolve ambiguities automatically. When multiple input metabolites correspond to the same unique identifier, or vice versa, these cases are flagged as ambiguities and described as 'Partial match' in the output. Users are then required to manually review and resolve these ambiguities to ensure data integrity. For illustration, we provide examples of three types of ambiguities, encountered in the PD study mapping procedure using the MetaNetX conversion table.

**One metabolite of the metabolomic annotation profile maps onto two distinct metabolites of the GSMN (Fig. S1)** The metabolite *3hivac* detected in the metabolomic dataset is mapped onto the GSMN; however, it is classified as an ambiguous mapping because another metabolite present in the Recon3_3 GSMN, namely *CE2028*, shares metadata. *3hivac* is identified directly in the metabolomic data via the VMH identifier. An InChI is also provided as additional metadata for this metabolite in the annotation profile; *3hivac* and the InChI are tested independently during the mapping process, although they are associated with the same entry in the annotation profiles. Thus, on the one hand, the VMH identifier is directly mapped to the IDs in the GSMN metadata (fig S1). On the other hand, the tested InChI has been mapped to the metabolic network for *3hivac* and *CE2028*, as this InChI is present in the metadata for both metabolites in the metabolic network. The two entries are therefore combined in the output and tagged as ambiguous mapping.

**One metabolite of the metabolomic profiles maps onto two identifiers in the conversion table (Fig. S2)** Such an ambiguity case occurred for metabolite *crn* that matched two MetaNetX compounds: *MNXM1105737* and *MNXM1105736. crn* is present in the metadata of both, representing (R) and (S)-carnitine. To handle this ambiguity, MetaNetMap will add both MetaNetX identifiers in the "partial match" column of the output; "MNXM1105736 _AND_ MNXM1105737" to warn the user that manual curation might be required.

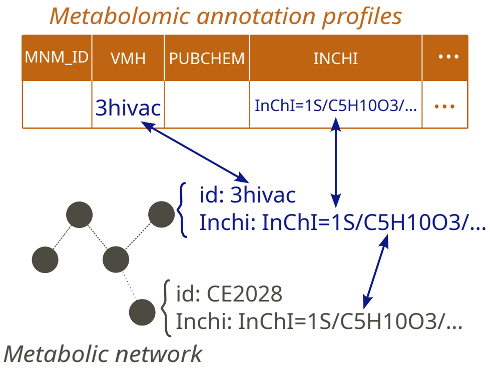


Figure S1: **Overview of a use case where one annotated metabolite maps onto two distinct compounds in the metabolic network**

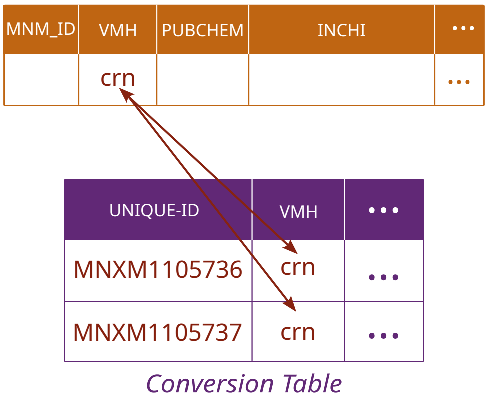


Figure S2: **Overview of a use case where one annotated metabolite maps to two distinct compounds in the conversion table**

**Two distinct metabolites of the metabolomic profiles map onto the same metabolite of the GSMN** We will present two examples from the PD case to illustrate this type of ambiguity.

The first case, depicted in Fig. S3, involves the MNM_IDs *MNM840* and *MNM845*, which end up being combined in the output. In the annotation profile, *MNM840* represents the metabolomic entry related to *PUBCHEM:11811*, while *MNM845* is associated with the VMH identifier *hLkynr*. Both *PUBCHEM:11811* and *hLkynr* therefore represent distinct entries in the metabolomic profile. However, the same InChI is associated with both entries. Thus, if a match is performed by MetaNetMap using InChI information, an ambiguity arises for both *MNM840* and *MNM845*: the two entries cannot be distinguished as distinct metabolites throughout the mapping. Interestingly, in the GSMN, the metabolite with the identifier hLkynr carries both a PubChem ID and an InChI as specified in the annotation file. Consequently, even if MNM840 or MNM845 had lacked the InChI annotation, an ambiguity would still have arisen from the GSMN point of view.

Another ambiguous case, depicted in Fig. S4, involves *MNM115* and *MNM132*, which are associated with the VMH compounds *CE2516* and *dlnlcg*, respectively. *CE2516* metabolomic entry is linked to two MetaNetX identifiers; *MNXM1370278* as a VMH identifier and *MNXM1370277* as a BiGG identifier. MNXM1370278 is additionally associated to a BiGG identifier: *dlnlcg* that is also the VMH identifier of *MNM132*. As a result, these two VMH identifiers from metabolomics collapse into a single metabolite, since MetaNetMap considers both of them indistinguishable during mapping. *MNM115* and *MNM132* are therefore merged in the output and flagged as a partial match, allowing the user to manually resolve this ambiguity.

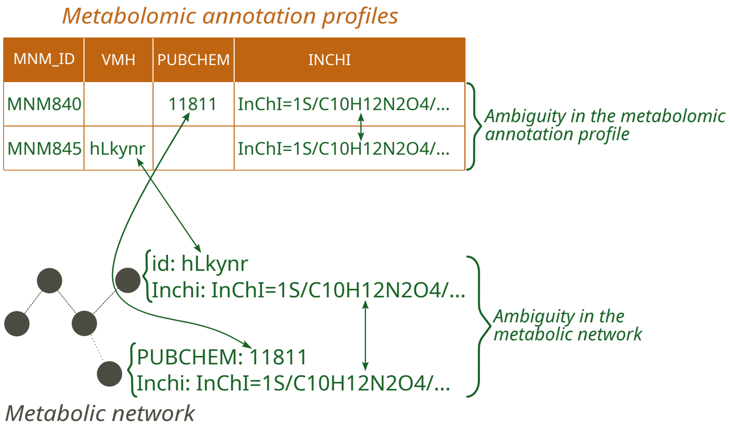


Figure S3: **Overview of a use case where two different annotated metabolites map onto two distinct compounds in the metabolic network, hence being merged in the output**

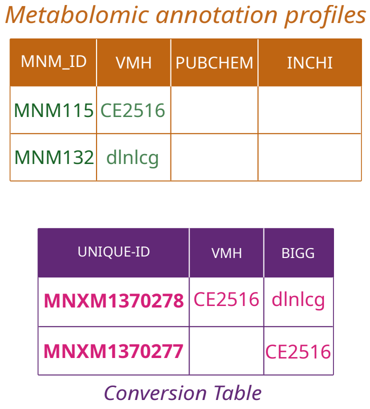


Figure S4: **Overview of a use case where two different annotated metabolites maps onto two distinct compounds in the conversion table, hence being merged in the output**

### 2 Supplementary methods on applications

#### 2.1 Data

Metabolomic profiles from the Parkinson's disease study were shared by the authors, and Recon3_3 GSMN was obtained from GitHub [2].
We downloaded metabolomic profiles for the mouse interventional study from Metabolights [2] project ID: MTBLS5608 [3] and used the same GSMN as the one for Parkinson's disease's applicaiton. To construct the reference databases (conversion tables), we used the compounds.dat file provided with MetaCyc version 29.0 and the chem_xref.tsv and chem_prop.tsv files from MetaNetX version 4.5.

#### 2.2 GSMN conversion

The Recon3_3 GSMN was originally in MATLAB (.dat) format; we converted it to SBML format to ensure compatibility with MetaNetMap. To achieve this, we used COBRApy [3] (v0.29.1) and ran the `cobra.io.load_matlab_model()` function to load the .dat file, followed by the `cobra.io.write_sbml_model()` function to export it in SBML format.

#### 2.3 Venn diagrams

We used the online tool InteractivVenn [4] to generate Venn diagrams and the associated dataset for the initial set and the set of interactions.

[2] https://github.com/opencobra/COBRA.papers/2024_metaPD
[3] https://www.ebi.ac.uk/metabolights/MTBLS5608

#### 2.4 Data preprocessing for application cases

The annotation profile from the PD study was extracted from Table 3 of 'Supplementary_Table1-8.xlsx' that was shared by the authors, and the header along with non-relevant columns were removed to produce a simplified table. This annotation profile consisted of two main columns: 'All', renamed 'COMMON-NAME' and containing both VMH identifiers and some 'PubChem' identifiers, and a separate column for 'InChI'. When diverse identifiers such as these are combined in a single column, the likelihood of successful matching is reduced, as numeric values and character strings are mixed. In contrast, standard practice promote the handling of numeric identifiers with appropriate prefixes, such as those corresponding to PubChem or ChEBI. Therefore, VMH and PubChem identifiers were separated as preprocessing.

The only preprocessing of metabolomic data applied to the mouse interventional study was renaming column names for compatibility with MetaNetMap.

### 3 Supplementary tables and figures

#### 3.1 Newly identified metabolites in the PD study

Tables S1 and S2 below list the additional metabolites that MetaNetMap could map onto the human metabolic network in the PD study, using MetaCyc and MetaNetX as conversion tables respectively. Identifiers in the first column refer to metabolomic entries, and the second column illustrates the identifiers that supported matches. Most of these identifiers were PubChem and InChI.

Rows highlighted in yellow indicate ambiguous cases, where the InChI identifier matches two distinct metabolites in the GSMN; the relevant identifiers are provided in the 'Ambiguities' column. Duplication of InChI or metabolite names in the metabolic network is responsible for these double matching cases. Rows highlighted in green indicate new metabolite matches identified solely using the MetaNetX database.

Certain ambiguities are specific to the conversion table used. For example, the metabolites *i* [4] and *ile_L* [5] share the same synonym in MetaCyc. These cases were not included in the table below, as they do not represent novel matches relative to the original study, but have nevertheless been documented in the output file.

[4]https://metacyc.org/META/NEW-IMAGE?object=I

[5]https://metacyc.org/compound?orgid=META&id=ILE

| New match | Match via | Ambiguities |
|---|---|---|
| 137tmurica1 | PUBCHEM – INCHI | |
| 17dmurt1 | PUBCHEM – INCHI | |
| 1mxnt | PUBCHEM – INCHI | |
| CE2028 | INCHI | 3hivac |
| 4hpro_LT | PUBCHEM – INCHI | |
| 5acam6am3mura | PUBCHEM – INCHI | |
| 5acam6fam3mura | PUBCHEM – INCHI | |
| 7mxth | PUBCHEM – INCHI | |
| appnn | PUBCHEM – INCHI | |
| CE2049 | PUBCHEM | |
| M03162 | INCHI | CE4888 |
| neuromelanin | INCHI | CE4888 |
| cfn1 | PUBCHEM – INCHI | |
| elaid | INCHI | ocdcea |
| HC02193 | PUBCHEM | |
| hexdeceeth | INCHI | pmeth |
| ocdcya | INCHI | lnlc |
| M00199 | INCHI | citr_L |
| M02146 | INCHI | ile_L |
| M01707 (dma) | | |
| phacgly | NAME | M02723 |
| pheacgly | NAME | M02723 |
| pcholn28_hs | NAME | |
| pxthn1 | PUBCHEM – INCHI | |
| slfcys | PUBCHEM | |
| theobromine1 | PUBCHEM – INCHI | |
| theophylline1 | PUBCHEM – INCHI | |

Table S1: **Newly identified metabolites using MetaNetMap (MetaCyc database) on the PD study**
*'New match'* column describe metabolomic entries that were mapped onto the metabolic network by MetaNetMap but were not mapped in the original study. *'Match via'* column indicates the identifiers that supported the match. *'Ambiguities'* column and yellow rows represent ambiguities between the newly identified metabolite in this column and a metabolite that was previously identified in the study. There were partial matches due to non unique InChI in the metabolic network.

| New match | Match via | Ambiguities |
|---|---|---|
| 137tmurica1 | PUBCHEM – INCHI | |
| 17dmurt1 | PUBCHEM – INCHI | |
| 1mxnt | PUBCHEM – INCHI | |
| CE2028 | INCHI | 3hivac |
| 4hpro_LT | PUBCHEM – INCHI | |
| 5acam6am3mura | PUBCHEM – INCHI | |
| 5acam6fam3mura | PUBCHEM – INCHI | |
| 7mxth | INCHI | |
| appnn | PUBCHEM – INCHI | |
| CE2049 | PUBCHEM | |
| M03162 | INCHI | CE4888 |
| neuromelanin | INCHI | CE4888 |
| cfn1 | PUBCHEM – INCHI | |
| elaid | INCHI | ocdcea |
| HC02193 | PUBCHEM | |
| hexdeceeth | INCHI | pmeth |
| ocdcya | INCHI | lnlc |
| M00199 | INCHI | citr_L |
| M02146 | INCHI | ile_L |
| M01707 (dma) | | |
| phacgly | NAME | M02723 |
| pheacgly | NAME | M02723 |
| pcholn28_hs | NAME | |
| pxthn1 | PUBCHEM – INCHI | |
| slfcys | PUBCHEM | |
| theobromine1 | PUBCHEM – INCHI | |
| theophylline1 | PUBCHEM – INCHI | |
| 2hiv | INCHI | |
| M00494 | INCHI–NAME | |
| M01051 | INCHI | |
| magpalm_hs | INCHI | |
| pchollinl_hs | INCHI | |
| pcholpalme_hs | INCHI | |

Table S2: **Newly identified metabolites using MetaNetMap (MetaNetX) on the PD study** *'New match'* column describe metabolomic entries that were mapped onto the metabolic network by MetaNetMap but were not mapped in the original study. *'Match via'* column indicates the identifiers that supported the match. *'Ambiguities'* column and yellow rows represent ambiguities between the newly identified metabolite in this column and a metabolite that was previously identified in the study. There were partial matches due to non unique InChI in the metabolic network.

#### 3.2 Comparison of mapping strategies

The following three Venn diagrams compare the sets of metabolomic entries mapped by distinct strategies:

- The PD application case using the original study using MetaNetMap with MetaCyc and MetaNetMap with MetaNetX.
- The mouse interventional study using MetaNetMap with MetaCyc and MetaNetMap with MetaNetX, both using the partial match option.

Counts consider all mapped metabolites, including ambiguities.

Figure S5 relates to the PD study. MetaNetMap retrieved all metabolites that were matched in the original study as well as additional ones that are database specific. MetaNetMap with MetaNetX generated the most matches.

Figure S6 compares the four strategies applied to the mouse interventional case study. MetaNetMap was run with MetaCyc and MetaNetX and with or without the partial matching option enabled. In this case, MetaCyc supports the most matches.

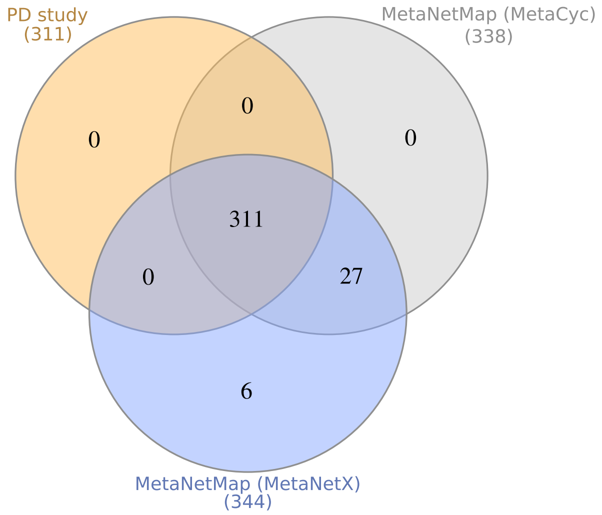


Figure S5: **Venn diagram of metabolite mapping for the PD case study vs. MetaNetMap (MetaNetX) vs. MetaNetMap (MetaCyc)**

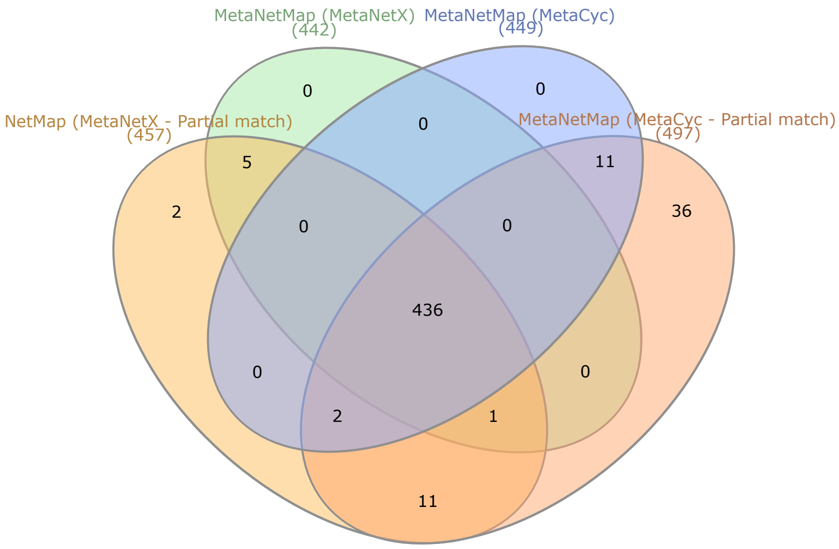


Figure S6: **Venn diagram of metabolite mapping for the mouse interventional study with MetaNetMap using MetaCyc or MetaNetX as conversion databases, and with or without partial match options**

[2] Ozgur Yurekten, Thomas Payne, Noemi Tejera, Felix Xavier Amaladoss, Callum Martin, Mark Williams, and Claire O'Donovan. Metabolights: open data repository for metabolomics. *Nucleic Acids Research*, 52(D1):D640–D646, 11 2023.

[3] Ali Ebrahim, Joshua A. Lerman, Bernhard O. Palsson, and Daniel R. Hyduke. COBRApy: COnstraints-based reconstruction and analysis for python. 7(1):74, 2013.

[4] Henry Heberle, Gabriela Vaz Meirelles, Felipe R da Silva, Guilherme P Telles, and Rosane Minghim. Interactivenn: a web-based tool for the analysis of sets through venn diagrams. *BMC bioinformatics*, 16(1):169, 2015.
